# Genetics and Environment Distinctively Shape the Human Immune Cell Epigenome

**DOI:** 10.1101/2023.06.29.546792

**Authors:** Wenliang Wang, Manoj Hariharan, Wubin Ding, Anna Bartlett, Cesar Barragan, Rosa Castanon, Vince Rothenberg, Haili Song, Joseph Nery, Andrew Aldridge, Jordan Altshul, Mia Kenworthy, Hanqing Liu, Wei Tian, Jingtian Zhou, Qiurui Zeng, Huaming Chen, Bei Wei, Irem B. Gündüz, Todd Norell, Timothy J Broderick, Micah T. McClain, Lisa L. Satterwhite, Thomas W. Burke, Elizabeth A. Petzold, Xiling Shen, Christopher W. Woods, Vance G. Fowler, Felicia Ruffin, Parinya Panuwet, Dana B. Barr, Jennifer L. Beare, Anthony K. Smith, Rachel R. Spurbeck, Sindhu Vangeti, Irene Ramos, German Nudelman, Stuart C. Sealfon, Flora Castellino, Anna Maria Walley, Thomas Evans, Fabian Müller, William J. Greenleaf, Joseph R. Ecker

**Author notes:** Equal contribution (co-first) (Wenliang Wang, Manoj Hariharan, Wubin Ding).

## Abstract

The epigenomic landscape of human immune cells is dynamically shaped by both genetic factors and environmental exposures. However, the relative contributions of these elements are still not fully understood. In this study, we employed single-nucleus methylation sequencing and ATAC-seq to systematically explore how pathogen and chemical exposures, along with genetic variation, influence the immune cell epigenome. We identified distinct exposure-associated differentially methylated regions (eDMRs) corresponding to each exposure, revealing how environmental factors remodel the methylome, alter immune cell states, and affect transcription factor binding. Furthermore, we observed a significant correlation between changes in DNA methylation and chromatin accessibility, underscoring the coordinated response of the epigenome. We also uncovered genotype-associated DMRs (gDMRs), demonstrating that while eDMRs are enriched in regulatory regions, gDMRs are preferentially located in gene body marks, suggesting that exposures and genetic factors exert differential regulatory control. Notably, disease-associated SNPs were frequently colocalized with meQTLs, providing new cell-type-specific insights into the genetic basis of disease. Our findings underscore the intricate interplay between genetic and environmental factors in sculpting the immune cell epigenome, offering a deeper understanding of how immune cell function is regulated in health and disease.

## Introduction

The debate between nature and nurture is a long-standing discussion in both biology and society. It centers around the relative impact of genetic inheritance (nature) versus environmental factors (nurture) on human development. In the field of epigenetics, the dividing line between an inherited and an acquired feature remains unclear ^1,2^. While inherited epigenetic marks are passed down through generations, acquired features arise from environmental influences and can alter gene expression without changing the underlying DNA sequence. Recent studies found that acquired epigenetic features can also be inherited ^3^. Understanding the contributions of these two sources of epigenetic variation is crucial for comprehending how genes and environments together shape biological outcomes. This complexity highlights the need for ongoing research to elucidate how inherited and environmental factors interact to affect the epigenome and further influence various aspects of health and disease.

The interplay between genetic predispositions and environmental factors shapes biological outcomes^4^. Monozygotic twin studies have enhanced our understanding of how genetic, environmental, and stochastic factors influence epigenetics ^5–8^. DNA Methylation as an important layer of epigenome, has been reported to have inheritance between generations ^3,9,10^. Previous twin studies based on bulk tissues have estimated that the average heritability of methylation levels at cytosine-guanine dinucleotides (CpGs) across the genome ranges from 5% to 19% in different tissues ^11–14^. Following studies on methylation quantitative trait locus (meQTL) studies have also revealed the association between genetic variations and methylation status of individual CpG sites ^15–20^. However, these studies use bulk tissues or whole blood, and most of them used Illumina Methylation EPIC array to profile the methylome. A comprehensive genome-wide and cell-type-specific relationship between genetic variations and methylation has yet to be fully elucidated.

The exposome, which includes the entirety of exposures—such as chemical, microbiological, physical, medicinal, and diet—that an individual encounters throughout their life, has the potential to influence the epigenome. The epigenome has long been recognized as a crucial intermediary influenced by environmental factors, a concept first highlighted by studies showing its role in regulating plant flowering in response to environmental cues ^21,22^ and monozygotic twins in human ^7^. The theme of *exposome and epigenetics* generates excitement because environmental exposures are increasingly linked to phenotypic changes and diseases through mechanisms involving DNA methylation and chromatin alterations ^23–25^. Human studies of how individual exposure can remodel the epigenome have been a focal point in recent single-cell research. For example, Aracena et al. ^23^ have revealed chromatin accessibility changes associated with influenza virus, and numerous coronavirus disease 2019 (COVID-19) studies have discussed the role of Severe acute respiratory syndrome coronavirus 2 (SARS-CoV-2) infection in reshaping the epigenome ^24^. However, these studies typically focus on chromatin accessibility changes and often investigate each exposure separately. Our understanding of how much the exposome will reshape the comprehensive epigenome in each cell type is still limited. Furthermore, as Feil et al. stated more than ten years ago, the relative contributions of extrinsic (environmental) factors and intrinsic factors to stochastic changes remain largely unknown ^25^. Additionally, the precise mechanisms by which genetics and environmental factors (G × E) interact to regulate gene expression and influence human health and disease remain an area of active research ^2^.

To dissect the contributions of the exposome and genetics to human immune cell epigenomes in a cell-type-specific manner, we obtained 171 peripheral blood mononuclear cell (PBMC) samples from 110 individuals. These individuals were either exposed or not exposed to various pathogens (Human immunodeficiency virus 1 [HIV-1], Influenza A virus [IAV], Methicillin-resistant *Staphylococcus aureus* [MRSA], Methicillin-sensitive *Staphylococcus aureus* [MSSA], Anthrax Vaccine [BA], SARS-CoV-2) and chemicals (Organophosphate [OP]). We performed Fluorescence-activated Cell Sorting (FACS) on major immune cells (B cell, Monocyte, NK cell, CD8 memory T cell [Tc-Mem], CD8 Naive T cell [Tc-Naive], CD4 memory T cell [Th-Mem], and CD4 naive T cell [Th-Naive]). Further, we performed single-nucleus methylation sequencing (snmC-seq2) ^26^ on these samples. Additionally, we conducted single-nucleus ATAC-seq ^27^ on the HIV-1 samples to further explore chromatin accessibility. We built a comprehensive catalog of cell type-specific epigenomic alterations by each individual exposure (eDMRs) and dissected the methylome that is determined by genetic variations (gDMRs) and affected by exposome. eDMRs and gDMRs are enriched in different chromatin regions, with eDMRs enriched at enhancer marks, while gDMRs enriched at gene body marks, indicating genetics and exposures regulate gene expression through different mechanisms. We found that each exposure can uniquely alter the epigenome and some exposures can perturb the enhancers which are probably bound by master transcription factors. Comparison of HIV-1-associated changes in methylation and chromatin accessibility showed significant correlations. We further found substantial colocalization of GWAS SNPs with the meQTLs, which provides cell type-specific insight on these disease SNPs. The human population immune cell atlas and the catalog of exposures and genetics-associated epigenomic features will be excellent resources for future mechanistic research on both human infectious and genetic diseases.

## Results

### An exposures-driven single-cell epigenomic atlas of human immune cells

We collected 171 PBMC samples from 110 individuals who were exposed or not exposed to seven major exposures (Figure 1, Table S1). HIV-1 exposed samples were collected from iPrEx cohort, a Phase III clinical trial, which was designed to assess the efficacy of pre-exposure prophylaxis (PrEP) for HIV-1 prevention. We analyzed PBMC samples from nine donors at three distinct time points: approximately 200 days before HIV-1 positivity (pre), the day of HIV-1 diagnosis (acu), and approximately 200 days after initiating treatment (cro). In IVA exposure, the BARDA-Vaccitech FLU010 study evaluated the VTP-100 vaccine against the H3N2 influenza virus strain. We studied pre- and 28 days post-challenge (with live H3N2 influenza virus) PBMC samples from 18 donors who received the placebo vaccine. We also analyzed PBMC samples from donors who were exposed to SARS-CoV-2 and had severe or non-severe COVID-19 symptoms.

**Figure 1.**
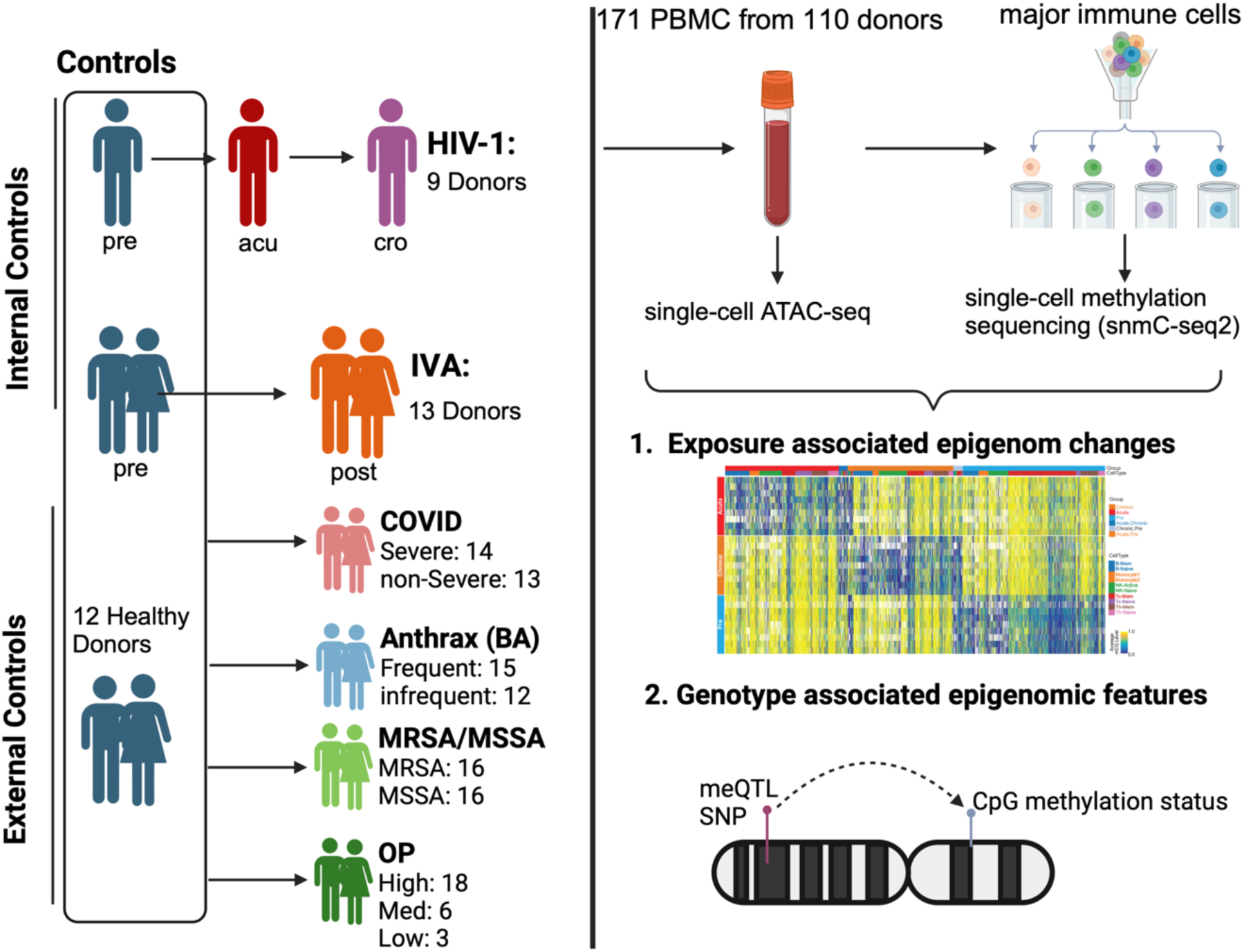
Overview of the study. For HIV-1 and IVA, we have internal control samples which are from the same set of donors before infection, and collected samples from them after exposure. We also collected PBMC from 12 healthy donors as external controls. For exposures without internal controls (COVID, Anthrax Vaccine, MRSA/MSSA and OP), all the healthy samples were used as control. We performed single-nucleus ATAC-seq and single-nucleus methylation sequencing on the PBMCs and identified the exposure associated differentially methylated regions (eDMRs) associated with exposures and genotypes. We also identified the genotype-associated DMRs (gDMRs) using this dataset.

For bacterial exposure, PBMC from 19 patients who tested positive to either MRSA or MSSA were analyzed, with a total of 27 samples. We also obtained PBMC samples from 27 vaccinated subjects who handled *Bacillus anthracis* in a controlled Biosafety Level 3 (BSL3) facility while wearing appropriate PPE. These individuals were trained scientists working in a BSL3 laboratory who had received either BioThrax®, an inactivated, acellular vaccine primarily containing the non-pathogenic protective antigen (PA) protein, or the Anthrax Vaccine Adsorbed as a part of the safety protocols of the facility. Organophosphates (OP) are a class of pesticides that are known to have a severe impact on the dopaminergic and serotonergic systems. A common form of this pesticide (Chlorpyrifos) has been used widely in the US. As part of this project, samples were collected from farm workers and residents around the farms. By tracking the levels of TCPY over a four-month period, samples from 27 donors were classified as exposed to high, moderate, or low levels of OP.

PBMCs from these samples were FAC-sorted into major immune cell types (Figure S1) and processed through a single-nucleus methylation sequencing pipeline as previously described ^26^. 96 cells for each sample and cell type were sorted to ensure enough coverage for each cell type. To control for potential batch effects, all cell types were sorted on the same 384-well plate (Figure S1). Additionally, we performed single-nucleus ATAC-seq on unsorted PBMCs from all three-time points from 4 HIV-1 donors. The analysis of the single-nucleus methylation data was performed using our in-house pipelines, as previously described^26^. After filtering out low-quality cells, 104,000 cells were clustered using average CG methylation level in 5 Kb bins across the autosomes. The single-nucleus ATAC-seq data from HIV-1 were integrated with methylation data and cell type labels were transferred.

### Heterogeneity in Methylation Profiles Reveals Exposure-Specific Immune Cell Clusters

To investigate whether the exposome globally affects the methylome of each immune cell type, we performed within-cell-type clustering for the major sorted immune cell types. This analysis revealed heterogeneity in methylation profiles, resulting in more than ten distinct clusters for each cell type. We observed significant bias of cells from each exposure in each cell type. Of note, HIV-1, SARS-CoV-2 and MRSA/MSSA exposures seem to have unique monocyte, CD4 and CD8 naive T cell profiles. (Figure 2A, Figure S2). This is unlikely to be a batch effect as different cell types from the same sample are processed on the same 384-well plate (Figure S3C). The proportion of immune cells from each exposure in these clusters varied significantly (Figure 2B). For example, we observed a cluster of monocytes enriched in both severe and non-severe COVID samples. Considering the biased distribution among the clusters of different exposures might also be caused by heterogeneity between individuals, but not the specific exposures, we focused on the proportional change of HIV-1 and IAV samples, which were from the same donors before and after infections. While IAVsamples have comparable proportions in each cluster across the cell types, HIV-1 infection markedly changed the cell proportions among clusters, indicating that HIV-1 remodeled the global methylome and functional states of these immune cells, especially in NK cells, CD8 memory and naive T cells (Figure 2C).

**Figure 2.**
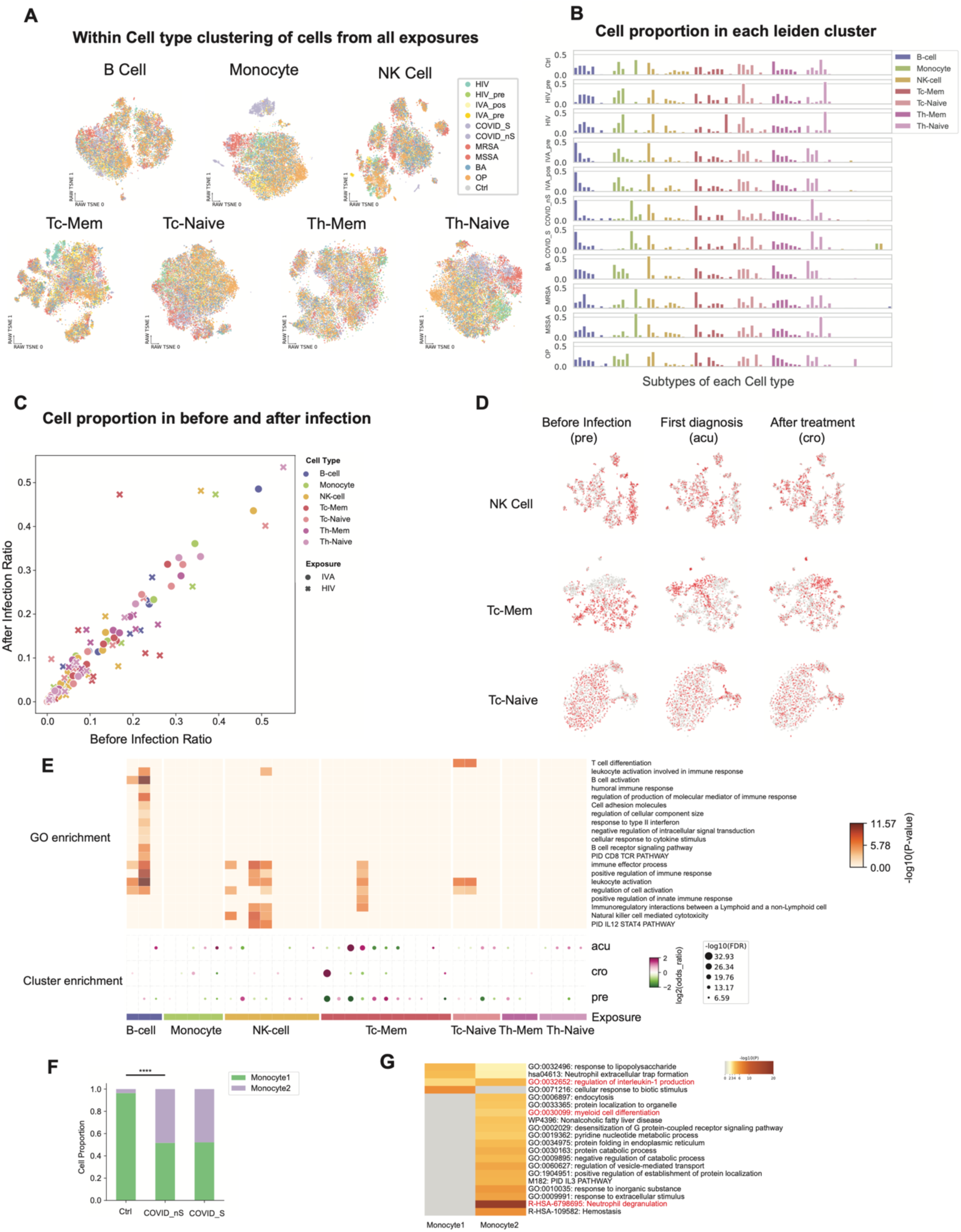
Within cell type changes are associated with each exposure. **A.** Uniform Manifold Approximation and Projection (UMAP) of cells in each cell type from FACS using single-nucleus methylation sequencing data. The cells are colored by exposures. HIV: exposure to HIV; HIV_pre: before HIV infection from the same donors; Flu_pos: after IVA infection; Flu_pre: before IVA infection from the same donors; COVID_S: severe COVID patient samples; COVID_nS: non-severe COVID patient samples; MRSA: samples exposed to MRSA; MSSA: samples exposed to MSSA; BA: samples from the donors that have taken Anthrax vaccine and work frequently or infrequently in a controlled BSL3 facility handling *Bacillus anthracis*; OP: samples from the donors that are exposed to OP. **B.** bar plots show the proportions of cells from each group and cell type in the Leiden clusters, colored by the FACS cell types. The x-axis are the Leiden clusters, and the y-axis shows the proportions of cells in each Leiden cluster in each group. **C.** Scatter plot shows the cell proportional changes before and after infection of HIV and IVA in each Leiden cluster. Dots are clusters with IVA exposures and crosses are HIV exposure. Color shows the FACS cell types. **D.** UMAP of cells from HIV exposure donors in the cell types that have the most cell proportion changes. The three rows are UMAP of cells from NK cell, Tc-Mem and Tc-Naive, and the columns are the cells from ‘pre’, ‘acu’ and ‘cro’ stages. Cells from the stage are shown in red, and cells from other stages are shown in gray. **E.** The dot-plot shows the enrichment of cells from the three HIV infection stages (‘pre’, ‘acu’, ‘cro’) in the Leiden clusters. The heatmap shows the GO enrichment of differentially methylated genes (DMGs) of the corresponding Leiden cluster. The dot plot and heatmap have the same x-axis. **F.** Two clusters of monocytes were identified. The bar plot shows the cell proportions of the two clusters of monocytes in controls, severe and non-severe COVID samples. Statistical tests were done using the Chi-Square test. (**** indicates P value < 1×10^−100^). **G.** GO enrichment of DMGs between the two clusters of Monocytes.

To further validate the changes in functional states of different immune cell types caused by HIV-1 infection, we performed within-cell-type clustering of HIV-1 samples. The clustering revealed that cells from the three stages of HIV-1 infection (pre, acu, and cro) were unevenly distributed among the clusters, with biased clusters exhibiting different global methylation levels (Figure 2D, Figure S3A). In NK cells, the cluster that is enriched in ‘pre’ stage cells has a higher global methylation level, which is likely in a naive state, suggesting a transition from naive to active NK cells after HIV-1 infection. Similarly, in CD8 naive T cells, the cluster that is hypomethylated is enriched in ‘acu’ and ‘cro’ stage cells, indicating these cells might be exiting the naive state to the effector state, though the naive T cell marker CCR7 is on the cell surface (FACS). To further dissect the identity of each subtype that is significantly differentially distributed among the three stages (FDR < 0.05, Fisher’s exact test), we identified differentially methylated genes (DMGs) in each cluster compared to all other clusters and performed functional enrichment of these genes. Immune cell activation and differentiation-related functions are enriched among these clusters (Figure 2E), suggesting that clusters that are biased to specific HIV-1 infection stages are related to a response to HIV-1 infection. While the marker genes in B cells and NK cells are driven by the global difference between naive states and memory/active states, the cluster in CD8 memory T cells that were specifically enriched in ‘acu’ stage cells are enriched in positive regulation of immune response. In other naive cell subtypes, where we didn’t observe an enrichment of immune activation functions, we observed some important immune genes are hypomethylated. For example, in a subtype of CD4 naive T cells which is depleted in ‘acu’ stage cells, TCF7 and LEF1 are hypomethylated, suggesting the acute stage CD4 naive T cells were exiting naive states.

While donor heterogeneity cannot be entirely excluded, the monocyte cluster uniquely enriched with COVID samples and depleted in controls and other exposures is likely associated with this specific exposure (Figure 2F). Almost half of the monocytes from both severe and non-severe COVID-19 samples are separated in these two clusters (Figure 2F), significantly more than the control samples (P=2.05e-237, Fisher’s exact test). In our sorting strategy, we specifically sorted CD14 high populations as monocytes (Figure S1A), so the two populations are not CD14 monocytes and CD16 (CD14 low) monocytes as identified by other single-cell studies. To further confirm the identity and function of the two clusters of monocytes, we identified the DMGs and DMRs between them. We identified 321 DMGs between ‘Monocyte1’ and ‘Monocyte2’, of which 262 and 59 genes are hypomethylated in ‘Monocyte2’ and ‘Monocyte1’, respectively (Figure S3B). Functional enrichment of these genes showed that genes hypomethylated in both monocyte clusters are enriched in pro-inflammation functions like ‘IL-18 signaling pathway’ and ‘regulation of interleukin-1 (IL-1) production’(Figure S4G). Both IL-18 and IL-1 are reported to be protective during murine coronavirus infection ^28^, while IL-1 has a pivotal role in the induction of cytokine storm due to uncontrolled immune responses in SARS-Cov2 infection ^29^. This suggests that both monocyte clusters exhibit pro-inflammatory signatures, though different genes are involved in the process. Moreover, besides IL-1 and IL-18 production-related functions, DMGs in ‘Monocyte2’ are also enriched in the functions of ‘phagocytosis’ and ‘endocytosis’ (Figure 2G), indicating the antigen presentation function of this cluster, which is specifically enriched in COVID-19 monocytes.

### Cell-Type-Specific Epigenomic Responses to Exposures

To comprehensively assess the impact of exposome on the epigenome in the immune cells in a cell-type-specific manner, we refined our cell types by integrating methylation information. Clustering with methylation profiles of all immune cells showed that B cells and NK cells can be separated into two clusters by global CG methylation level (Figure 3A). The higher methylated clusters were annotated as a naive state (B-Naive and NK-Naive), while the cluster with lower methylation level of B cells is annotated as memory B cells (B-Mem), and a lower methylated cluster of NK cells is defined as the active state (NK-Active)^30^. We observed some inconsistency between FACS and methylation profile clustering, with some naive T cells clustered together with memory T cells, indicating the transition of cell states of some naive T cells (Figure 3A). We labeled these cells still as naive T cells based on cell surface marker CCR7+ and CD45RA+. Some sorted NK cells (CD56+ or CD16+ and CD14-) also have a similar methylation profile as monocytes, which might be some CD16+ monocytes. These cells were not included in the downstream analysis. In the following analysis, we dissected the contributions of exposome and genetics to the epigenome in these nine immune cell types.

**Figure 3.**
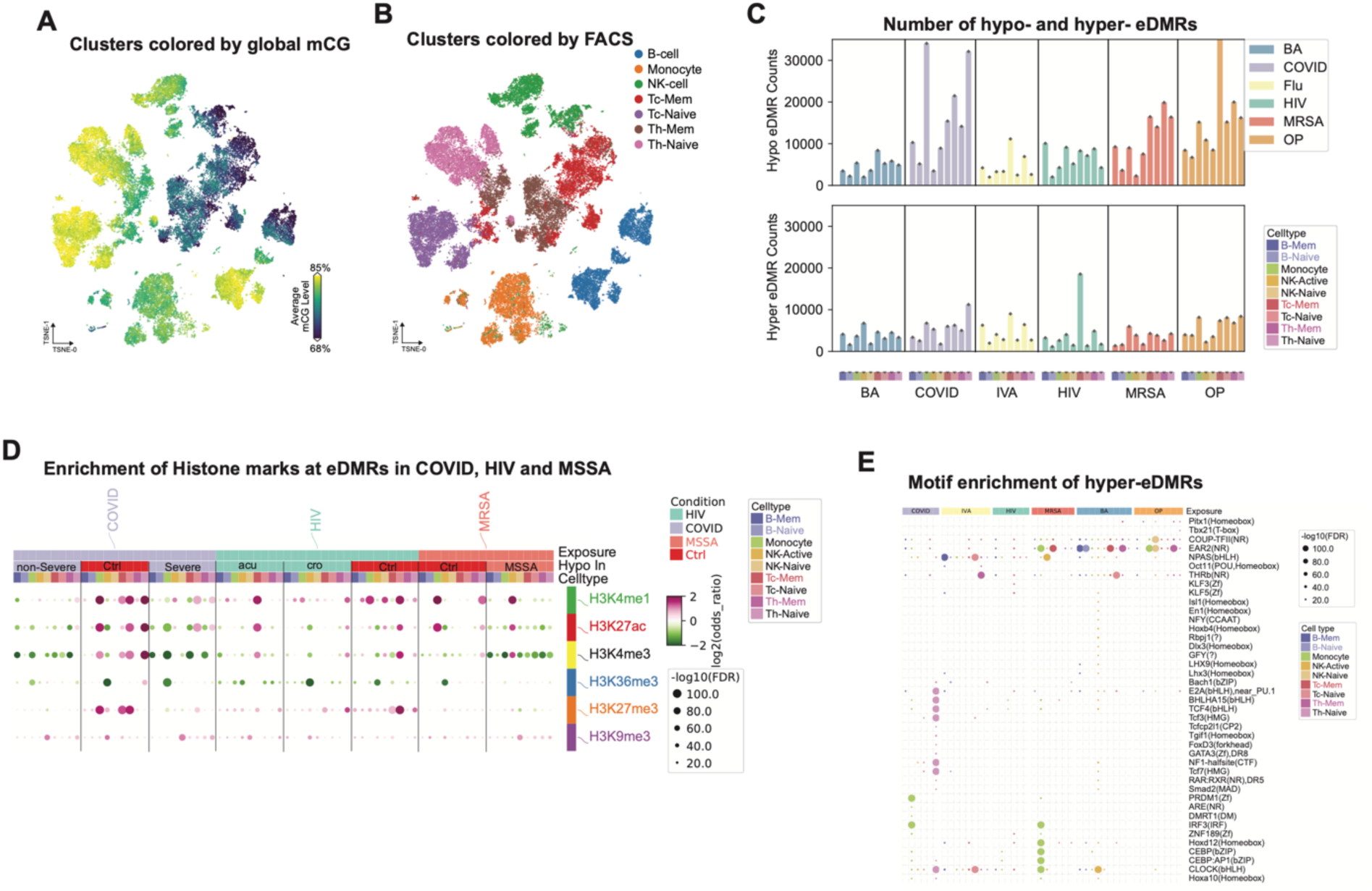
Identification of Exposure-associated DMRs (eDMRs) and their features. **A.** UMAP of all cells from all exposures using single-nucleus methylation profiles. Cells are colored by the global methylation level of each cell. **B.** UMAP of all cells from all exposures using single-nucleus methylation profiles. Cells are colored by the FACS cell types. **C.** Barplots shows the Hypo-(upper plot) and Hyper-(lower plot) methylated eDMR counts. Hypo means the eDMRs are hypo-methylated in exposures compared to controls. Hyper is the other way around. The colors show the exposures, and x-axis is cell type, which is colored and sorted in the same order for all exposures. **D.** Dotplot shows the enrichment of eDMRs from COVID, HIV and MSSA in histone modification peaks. Each column shows the hypo-eDMRs in that condition. Color of the dots shows the enrichment or depletion in the corresponding histone modification. **E.** Dotplot shows the motif enrichment of eDMRs from each exposure and cell type. The dot size indicates the P values of enrichment, and the color shows the cell type from which the eDMRs are.

We aimed to dissect the impact of different exposures on the epigenome in these cell types by identifying the exposure-dependent differentially methylated regions (eDMRs). Considering it’s difficult to control other exposures each individual might have experienced and the different genomes of these donors, we used internal controls for HIV-1 and IVA exposures, for which we have cells before and after exposure. We used a stringent pipeline to identify the eDMRs of other exposures (Methods), in which the external controls are used. We used three sets of controls (healthy donors, HIV-1 ‘pre,’ and Flu ‘pre’), and identified eDMRs that were significantly different between exposure samples and all control samples. These eDMRs were not significantly different between any two sets of controls, minimizing the contribution of individual genetic variation and baseline exposures to these eDMRs.

We identified 756,575 eDMRs across all exposures and cell types, with 517,698 and 238,877 eDMRs hypo- and hyper-methylated, respectively (Figure 3C). On average, each exposure and cell type exhibited approximately 10,000 eDMRs. SARS-CoV-2, organophosphates (OP), and MRSA/MSSA showed the most abundant eDMRs across the majority of the cell types, highlighting their pronounced impact on the epigenetic profiles (Figure 3C). Of note, the majority of eDMRs in each exposure and cell type are single CpG sites (Figure S4A), suggesting that exposure-associated DNA methylation alterations are often confined to individual CpG sites. These single-CpG methylation changes have been reported in aging, development, environmental and disease changes^31–35^, and a recent study using nanopore sequencing also reported that most CpG units are singletons ^36^.

To further characterize the chromatin state of cell-type-specific eDMRs, we examined the associated histone modification peaks from ENCODE^37^ in each cell type. We found that both hypo- and hyper-methylated eDMRs across all exposures are slightly enriched in heterochromatin regions marked by H3K9me3 (Figure 3D, S4A). Notably, COVID-19 hyper eDMRs in monocytes, naive T cells, and NK-naive cells showed significant enrichment in enhancer marks like H3K27ac and H3K4me1, as well as the promoter mark H3K4me3, across all immune cell types (Figure 3D). Similar patterns were observed with HIV-1, where hypo eDMRs in CD8 memory T cells and hyper eDMRs in CD8 naive T cells displayed enrichment in these active histone marks (Figure 3D). Furthermore, MSSA monocyte hyper eDMRs were enriched in H3K27ac, while NK-active cells from MSSA-infected individuals (but not MRSA-infected ones) showed enrichment in enhancer regions. Interestingly, naive T cells from severe COVID-19 patients had hypo eDMRs with greater enrichment in active histone marks than those from non-severe patients (Figure 3D). These eDMRs are generally depleted at CG islands, promoters and SINE elements but slightly enriched in DNA, LINE and LTR transposable elements (Figure S4B).

DNA methylation influences transcription factor (TF) binding to DNA ^38,39^, potentially altering gene expression. To further investigate which transcription factors (TF) might be perturbed at these eDMRs, we performed motif enrichment for the hypo and hyper eDMRs separately. To compare the differentially enriched transcription factor motifs among the exposures and cell types, we performed principal component analysis (PCA) on the enriched motif of hypo and hyper eDMRs separately and selected the TF motifs in the top 10 PCs. The TF motifs in hypermethylated eDMRs display distinct enrichment patterns across each cell type and exposure, underscoring their unique perturbation on the epigenetic landscape (Figure 3E). Specifically, we observed master TF motifs were enriched in COVID-19 monocytes and CD4 naive T cells. For example, PRDM1 (encoding BLIMP-1) in monocytes ^40^ and TCF family TFs (including TCF7) motifs in CD4 naive T cells are uniquely enriched in COVID-19 samples ^40,41^. Similarly, CEBP family TF motifs are specifically enriched in MRSA/MSSA monocytes ^42^. The enrichment of hyper eDMRs from these two exposures and cell types in H3K27ac-marked enhancers suggests that the binding of master transcription factors in the corresponding cell types may be affected, potentially perturbing their functions due to these exposures. These results suggest that methylomes remodeled by exposures might be able to inhibit the binding of master TFs, through which they change the function of the cells. On the other hand, hypo-methylated eDMRs across the exposures share many similarly enriched motifs (Figure S4C).

### Linking DNA Methylation and Chromatin Accessibility in HIV-1 Exposure

To investigate whether the DNA methylation changes observed with each exposure correspond to alterations in chromatin accessibility, we conducted single-nucleus ATAC-seq on four of the same donors previously analyzed for methylation in the context of HIV-1 exposure. Comparing exposure-associated changes in DNA methylation and chromatin accessibility will provide a comprehensive catalog of epigenomes remodeled by exposure and also further dissect the eDMRs for following mechanistic investigations.

We integrated single-nucleus DNA methylation data with single-nucleus ATAC-seq data using 5 kb bins on autosomes. We mapped the cells from single-cell ATAC-seq to methylation clusters and transferred the cell type labels using Canonical Correlation Analysis (CCA) (Figure 4A, 4B)(Stuart et al., 2019). To assess how much the two modalities are correlated, we calculated the genome-wide correlation between these two modalities. Specifically, we divided the genome into 5 kb bins and calculated the hypomethylation score using single-cell methylation data and the number of Tn5 insertions using single-nucleus ATAC data. We then calculated the correlation between these two measurements for each bin. We observed a strong correlation between the two modalities (methylation and open chromatin) across all cell types, with the highest correlation observed in monocytes (Figure S5). Interestingly, we detected a loss of methylation and an increase in accessibility after HIV-1 infection in memory CD8 T cells at the intron of DGKH (Figure 4C), a gene previously reported to exhibit differential methylation between elite controllers (individuals able to maintain undetectable viral loads for at least 12 months without antiretroviral therapy) and individuals receiving antiretroviral therapy (Frias et al., 2021). Although this region experienced a loss of chromatin accessibility, the methylation level remained unchanged between the “acute” and “chronic” stages (Figure 3E). When comparing all eDMRs with differentially accessible regions (DARs) from the two modalities, we count the eDMRs that have the same direction (hypo eDMR with gained accessibility in the same stage) DAR within 1 kb and found a considerable fraction of eDMRs are consistent with changes in chromatin accessibility (Figure 4D). The highest overlap (25.6%) between these two modalities was observed in ‘pre’ stage hypo eDMRs in CD8 naive T cells. Given the overall correlation of these two modalities in each cell type, the overlap between hypo eDMRs and gained peaks indicates a consistent change in DNA methylation and chromatin accessibility.

**Figure 4.**
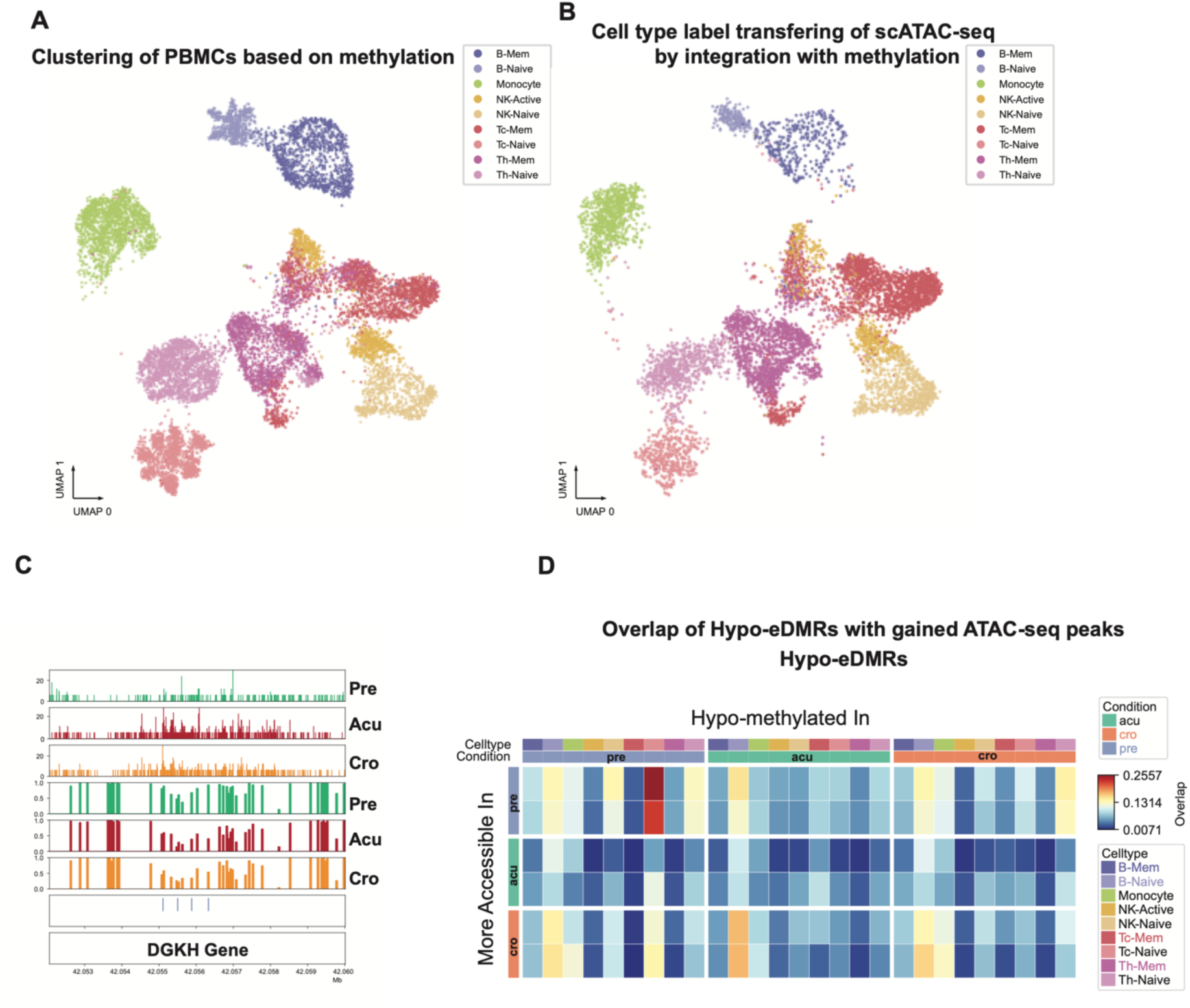
Consistent changes in DNA methylation and chromatin accessibility in HIV exposure. **A.** UMAP of cells from HIV exposure using single-nucleus methylation when integrating with single-nucleus ATAC-seq data. Color shows the cell types identified from FACS and DNA methylation. **B.** UMAP of cells from one HIV donor sample after integration with single-nucleus methylation data. The color shows the cell type labels transferred from DNA methylation data in integrating the two modalities. **C.** A genome browser view of a region at DGKH gene that has consistent changes after HIV exposure in DNA methylation and chromatin accessibility in CD8 memory T cells. The top three panels are normalized ATAC-seq reads, and the DNA methylation panels show the methylation levels in each bin at this locus. The eDMRs are shown in the blue bars. **D.** The overlap between hypo-eDMRs and gained ATAC-seq peaks in each condition in the corresponding cell types. The more accessible peaks are from pairwise comparisons, so each condition has two comparisons. The color of the heatmaps shows the proportion of overlaps between hypo-eDMRs and gained peaks.

### Genotype Associated DMRs Reveal the Influence of Genetics on Immune Cell Epigenomes

To dissect how much of the epigenome is determined by each individual’s genome, we used this single-cell epigenomic atlas to identify genotype-associated DMRs (gDMRs). We identified the single nucleotide polymorphism sites (SNPs) of each individual using the DNA methylation data with biscuit^43^, followed by genotype imputation with Minimac4 and filtering out SNPs that are not in dbsnp. To confirm the accuracy of the SNP calls using this strategy, we compared the SNPs from whole genome sequencing (WGS) and methylation data from a previous study from our lab, which used the same technology and platform. The results showed a strong agreement with WGS SNPs in a 10 Mb region, with low rates of false positives and false negatives (Figure S6A). To further verify the SNP accuracy from biscuit, we called the SNPs from our inhouse bulk m3C (a joint assay of DNA methylation and chromatin conformation) data from GM12878 cell line and compared the SNP calls with genotypes of NA12878. The SNPs from biscuit are quite accurate (Figure S6B). These results give us confidence to conduct genotype-associated epigenome analysis using SNPs derived from methylation data.

We first identified differentially methylated regions (DMRs) between individuals within each cell type in the CpG context and quantified the methylation level for each individual. DMRs overlapping with SNPs were excluded. Since the SNPs are derived from methylation data, those located in blacklist regions were also removed from downstream analysis. We then performed meQTL analysis as described (Methods).

After stringent filtering of the meQTL-DMR pairs, we got 234,600 gDMRs across all nine cell types, with 183,990 cis-correlated with SNP and 50,610 in trans. The number of cis gDMRs are comparable across different cell types except CD8 memory T cells, which have more gDMRs compared to other cell types (Figure 5A). The trans gDMRs are mostly identified in different T cell types (Figure 5A). To compare with the chromatin states of eDMRs, we did an enrichment analysis between these two sets of DMRs on different histone marks. In contrast to eDMRs at enhancer marks, gDMRs are predominantly enriched at gene body mark H3K36me3 peaks, especially in memory state lymphocytes (B cell, CD4 and CD8 T cells) (Figure 5B), while eDMRs are enriched at enhancer and promoter regions in naive lymphocytes. To further investigate the enrichment of chromatin loops between these two sets of DMRs, we did enrichment analysis of them at the loop anchors in different immune cells from ENCODE, which showed that gDMRs are more enriched at loops anchors in memory state lymphocytes and eDMRs are more associated with naive lymphocytes (Figure 5C). This indicates that eDMRs and gDMRs might regulate gene expression in different cell types through distinct mechanisms. This differential enrichment underscores the complexity of epigenetic control, with eDMRs and gDMRs contributing uniquely to the gene expression landscape depending on the cell type and the nature of the environmental or genetic input. Both eDMRs and gDMRs have similar genomic feature enrichment regarding genes and transposable elements (Figure S6C).

**Figure 5.**
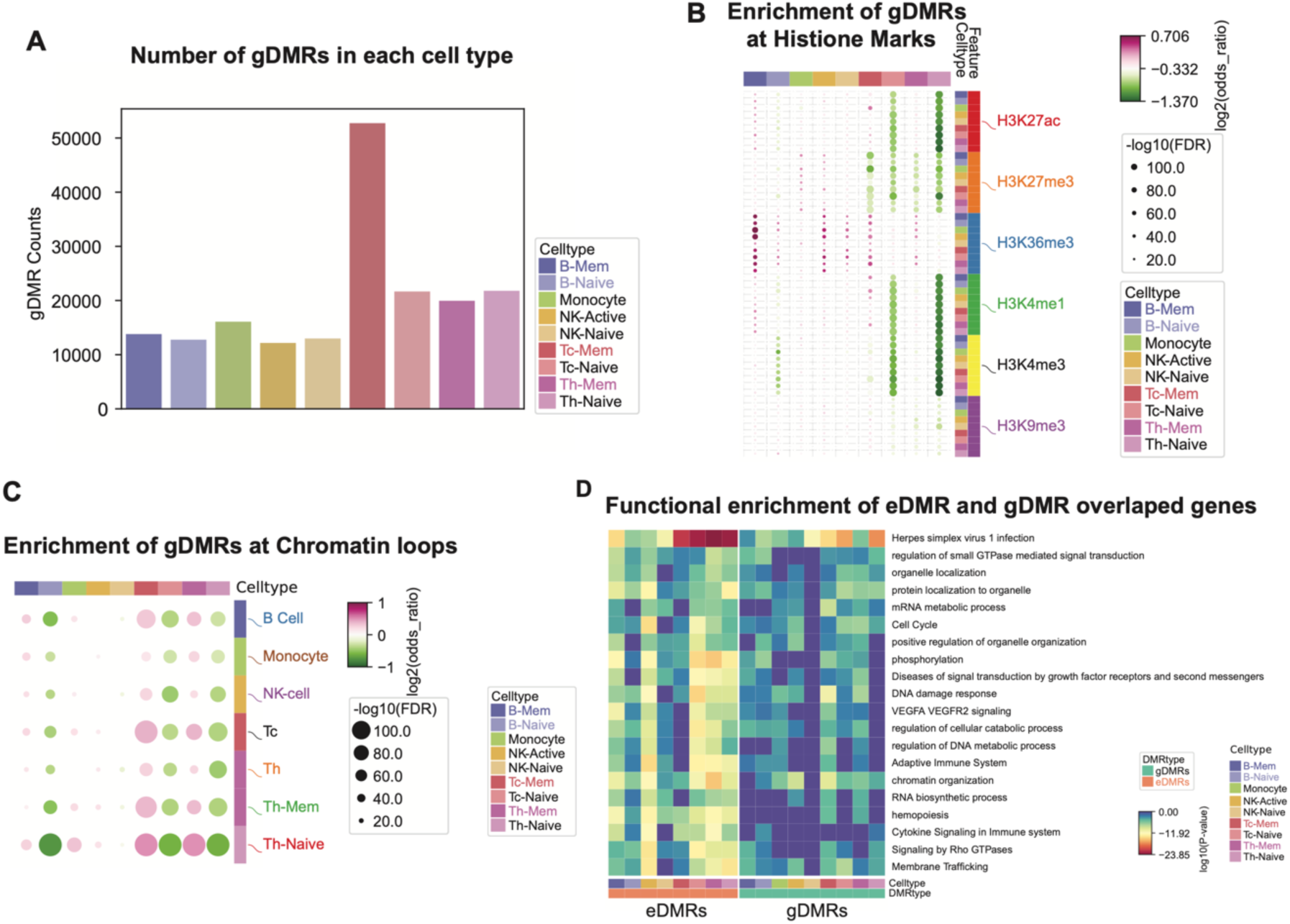
Identification of genotype-associated DMRs (gDMRs) and their features. **A.** barplot shows the counts of gDMRs in each cell type. The colors of the bars show the cell types. **B.** Dotplot shows the enrichment of gDMRs in histone modification peaks from each cell type. Each column is a cell type, and each row has one histone modification peak in each cell type. Color of the dots shows the enrichment of depletion of gDMRs in the corresponding histone modification. **C.** Dotplot shows the enrichment of gDMRs in chromatin loops. Each column is a cell type, and each row is the chromatin loop in each cell type. Color of the dots shows the enrichment or depletion of gDMRs in the corresponding chromatin loop. D. Go enrichment of the genes that are overlapped with eDMRs or gDMRs. Color of the heatmap shows the log10(P-value) of the enrichment. Results for eDMRs and gDMRs in all cell types are sorted in the same order.

To determine which genes might be regulated by eDMRs and gDMRs that overlap with H3K36me3 marks, we performed functional enrichment analysis on genes with DMRs overlapping H3K36me3 peaks. Both eDMRs and gDMRs were significantly enriched in housekeeping and immune-related functions, with more significant enrichment in eDMRs (Figure 5D). This suggests that eDMRs play a direct role in regulating the immune response to environmental exposures, whereas gDMRs in these immune cells are less involved. This distinction highlights the specific impact of environmental factors on immune-related gene regulation through epigenetic modifications.

Besides chromatin states, the enriched motifs also differ between eDMRs and gDMRs. While both hypo and hyper eDMRs are enriched mainly in immune-related TF motifs, gDMRs do not show enrichment of immune TF motifs (Figure S6D). Using the gDMRs as background in Homer, we found significant enrichment of RUNX and ETS family TF motifs, including PU.1, ETS1, and Fli1, in the eDMRs. In contrast, no significant motif enrichment except ETS motifs in CD8 memory T cells was observed in the gDMRs using eDMRs as background. These results suggest that compared with gDMRs, eDMRs are primarily enriched with key transcription factor binding sites in immune cells, playing a role in regulating gene expression through transcription factor binding. Meanwhile, gDMRs are more likely to be located on gene bodies, where they probably regulate gene expression in a different mechanism.

### Cell-Type-Specific Colocalization of gDMRs and GWAS SNPs Links to Immune Diseases

To investigate the association of gDMRs with human diseases and immune-related traits, we performed colocalization analysis between our meQTLs and GWAS SNPs linked to various traits. We identified many colocalized GWAS SNPs and meQTLs within these immune cell types, suggesting potential cell type-specific regulatory connections between methylation changes and genetic variants associated with immune functions. Enrichment analysis of GWAS SNPs within the meQTLs of each cell type revealed a predominant enrichment in CD8 naive T cells, as well as in CD4 memory and naive T cells (Figure S7A). This suggests that these specific immune cell types are particularly influenced by genetic variants associated with immune-related traits and diseases. For example, GWAS SNPs associated with “Alzheimer’s disease and Lewy body co-pathology” are enriched in CD8 naive T cells and CD4 memory T cells meSNPs that colocalize with the GWAS SNPs (Figure S7A). We further linked the gDMRs to specific diseases and phenotypes (Figure S7B) through colocalization analysis between meQTLs and phenotype-associated GWAS SNPs. The result showed that most gDMRs are only associated with one phenotype (Figure S7B).

This analysis enabled us to uncover cell-type-specific regulatory connections between gDMRs and various diseases and phenotypes, providing insights into the potential mechanisms by which GWAS SNPs influence the epigenome. For instance, SNPs associated with Addison’s disease showed colocalization with multiple meQTLs across various immune cell types in a cell-type-specific manner. Most of the GWAS SNPs only colocalize with meQTL in one cell type (Figure 6A), indicating a high degree of cell-type specificity in the regulatory effects of these GWAS SNPs on the epigenome. This suggests that the impact of genetic variants on DNA methylation, and consequently on gene regulation, can be highly specialized and confined to particular immune cell types. Notably, the SNP rs910320, associated with Addison’s disease, resides in the HLA locus and colocalized with a meQTL in CD8 naive T cells (Figures 6B, 6C), highlighting a potential epigenetic regulatory pathway specific to this cell type. This cell-type-specific colocalization information will greatly facilitate the mechanistic studies on the diseases.

**Figure 6.**
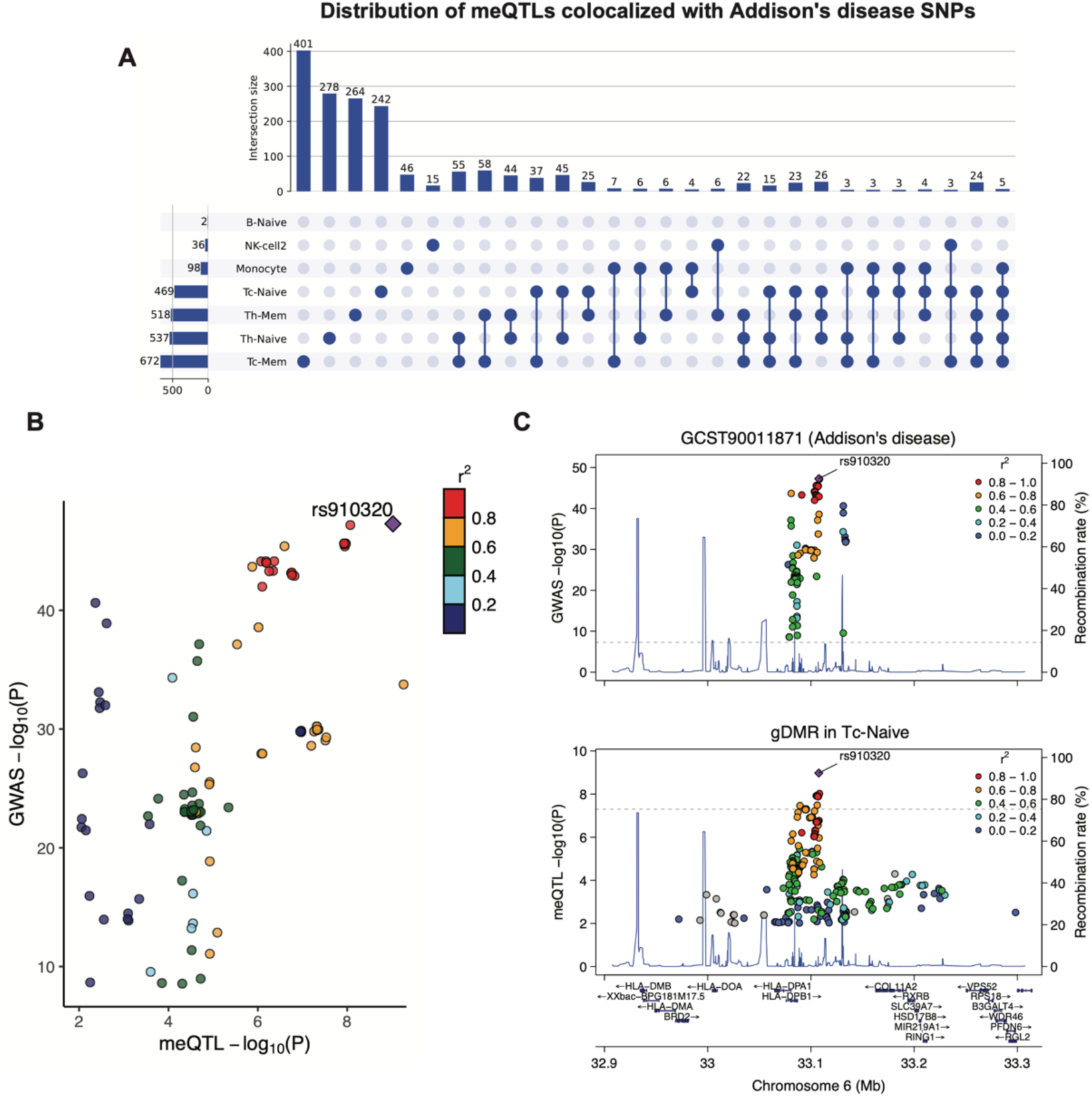
Colocalization of gDMRs with autoimmune-related disease (Addison’s disease) GWAS SNPs. **A.** Upset plot shows the number of meQTLs that are colocalized with GWAS SNPs in each cell type or multiple cell types. **B.** scatter plot shows the correlation of SNPs at the locus by the -log10(P-Value) associated with disease and gDMR. The r^2^ is the linkage disequilibrium with the leading SNP (rs910320). **C.** Genotype-disease association P values in the HLA locus for Addison’s disease GWASs (top panel) and meQTL signal in CD8 Naive T cell (bottom panel).

## Discussion

Our study delivers a comprehensive, exposure-driven atlas of human immune cells, revealing how genetic and environmental factors shape their epigenomes. We constructed an intricate epigenomic atlas using single-nucleus methylation sequencing and ATAC-seq, revealing the epigenomic features associated with different exposures. This comprehensive approach enabled us to identify exposure-specific DMRs (eDMRs) and genotype-associated DMRs (gDMRs), dissecting the roles of these two factors in shaping the epigenome of human immune cells. We also identified significant colocalization of meQTLs with GWAS disease-associated SNPs, uncovering the potential cell-type-specific epigenetic mechanisms of these SNPs.

Single-cell technologies have revolutionized our understanding of how individual cells’ transcriptomes and chromatin accessibilities respond to various exposures, such as smoking and infectious diseases ^7,44–46^. Our study primarily focuses on selected exposomes and their effect on DNA methylation, fills the gap of our understanding of how different exposures can alter the methylome of single cells. The eDMRs will serve as valuable resources for understanding the mechanisms underlying each exposure. Additionally, they have potential as biomarkers for identifying specific exposures, offering enhanced diagnostic and possibly therapeutic options. The unique subclusters associated with HIV-1 and SARS-CoV-2 exposures were not observed in other single-cell modalities ^47,48^, indicating distinct DNA methylation responses to these pathogens. Our approach, which combines FACS with single-cell methylation sequencing, enables us to investigate the heterogeneity within each cell type more effectively. The enrichment of specific exposure associated eDMRs in enhancers and promoters marks, which are mostly single-CpG changes, also deepens our understanding of these diseases.

Genetics and exposome have long been recognized to shape epigenomes ^4^. While previous studies have identified DNA methylation changes associated with genetic ^4,15–17^ or environmental exposures ^2,4,6^, our study uniquely examines both eDMRs and gDMRs within the same group of donors. This approach allows for a more reliable comparison of changes driven by these two factors. The differential enrichment of eDMRs and gDMRs on the chromatin indicates genetics and environments may regulate gene expression differently. Although we cannot fully disentangle the effects of genetics from different exposure histories in our donors, genetic factors exert a stronger regulatory influence on memory lymphocytes. In contrast, exposome regulation has a more pronounced effect on naive lymphocytes.

In summary, our findings highlight the complex contributions of genetic and environmental factors in shaping the epigenetic landscape of immune cells. This research enhances our understanding of how immune cell function is regulated by these factors and lays the groundwork for future studies exploring their combined effects on health and disease. The underlying mechanisms by which genetic factors and exposome modulate the effects of each exposure remain to be elucidated. The interplay between genetic and environmental factors in shaping the epigenome and influencing disease states remains to be fully uncovered. This gap in understanding highlights the need for further research to uncover how genetic predispositions interact with environmental factors to shape the epigenomic landscape, fully addressing the ‘nature and nurture’ question in human disease.

## Methods

### DATA GENERATION

#### Fluorescence-activated Cell Sorting of immune cell types

Cells were sorted into 384-well plates using fluorescence-activated cell sorting (FACS) based on their specific antibody labeling. The FACS antibody cocktail allowed for the identification of seven different immune cell types in blood (Figure S1). The sorted cell types included Naive helper T cells (CD3+, CD4+, CCR7+, CD45RA+), Memory helper T cells (CD3+, CD4+, CD45RA-), Naive cytotoxic T cells (CD3+, CD8+, CCR7+, CD45RA+), Memory cytotoxic T cells (CD3+, CD8+, CD45RA-), B cells (CD3-, CD19+), Monocytes (CD3-, CD19-, CD14+), NK cells (CD3-, CD19-, CD14-, CD16+, CD56+), and other cells (CD3-, CD19-, CD14-, CD16-, CD56-). The SONY Muti-Application Cell Sorter LE-MA900 Series was used to isolate single cells in 384-well PCR plates containing protein kinase. After cell sorting, the plates were spun down to capture the cells at the bottom of the well and then subjected to thermocycling at 50 ℃ for 20 minutes. The plates containing the DNA from the cells were subsequently stored at -20 ℃ or moved directly to library preparation.

#### snmC-seq2 Library preparation and Illumina sequencing

For library preparation, we followed the previously described methods for bisulfite conversion and library preparation in snmC-seq2 (Luo et al., 2017, 2018). The snmC-seq2 libraries generated from the isolated immune cells were sequenced using an Illumina Novaseq 6000 instrument with S4 flow cells in the 150-bp paired-end mode. Freedom EVOware v2.7 was utilized for library preparation, while Illumina MiSeq control software v3.1.0.13 and NovaSeq 6000 control software v1.6.0/Real-Time Analysis (RTA) v3.4.4 were employed for sequencing.

#### Single-nucleus ATAC-seq on HIV-1 donors

Single-nucleus ATAC-seq was performed as previously described ^49^, using the Chromium Next GEM Single Cell ATAC Library & Gel Bead Kit v1.1 (10x Genomics, 1000175) and the Chromium Next GEM Chip H (10x Genomics, 1000161) or Chromium Single Cell ATAC Library & Gel Bead Kit (10x Genomics, 1000110). Libraries were sequenced on the Illumina NovaSeq 6000 system (1.4 pM loading concentration, 50 × 8 × 16 × 49 bp read configuration), targeting an average of 25,000 reads per nucleus.

### QUANTIFICATION AND STATISTICAL ANALYSIS

#### Single-cell methylation data processing (alignment, QC)

For alignment and quality control (QC) of the single-cell methylation data, we employed the same mapping strategy used in our previous single-cell methylation projects in our lab (Liu et al., 2021). Specifically, we utilized our in-house mapping pipeline, YAP (https://hq-1.gitbook.io/mc/), for all the mapping-related analysis. The pipeline includes the following main steps: (1) demultiplexing FASTQ files into single cells, (2) reads-level QC, (3) mapping, (4) BAM file processing and QC, and (5) generation of the final molecular profile. Detailed descriptions of these steps for snmC-seq2 can be found in the work by Luo et al. (2018). All the reads were mapped to the human hg38 genome, and we calculated the methylcytosine counts and total cytosine counts for two sets of genomic regions in each cell after mapping.

We filtered out low-quality cells based on three metrics generated during mapping: mapping rate > 50%, final mC reads > 500,000, and global mCG > 0.5. Chromosomes X, Y, and M were excluded from the analysis, and the remaining genome was divided into 5 kb bins to create a cell-by-bin matrix. In this matrix, each bin was assigned a hypomethylation score (hypo-score) calculated from the p-values of a binomial test, which indicates the probability of hypomethylation of that bin. The matrix was further binarized for downstream analysis using a hypo-score cutoff of >= 0.95.

Hypo-score measures the likelihood of observing greater than *m* methylated reads under the assumption that methylation follows the binomial distribution with parameters *c* and p.

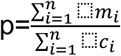

*m* is the observed number of methylated count for region *i*, *c* is the coverage (total count) covering region *i* and *n* is the total number of 5kb bin regions, p is the expected probability of methylation for this cell.

Let’s assume

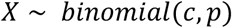

then for each 5 kb bin,

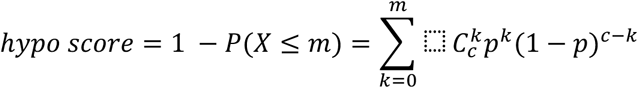

The calculation of hypo score was implemented in ALLCools (https://lhqing.github.io/ALLCools/intro.html) using scipy^50^

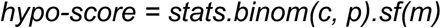

Bins covered by fewer than five cells and those with any absolute z-score(the number of cells with non-zero values) greater than 2 were filtered out. Additionally, we excluded bins that overlapped with the ENCODE blacklist using “bedtools intersect” (Dale, Pedersen, and Quinlan 2011; Quinlan and Hall 2010).

#### Unsupervised clustering

To perform unsupervised clustering, we utilized ALLCools (Liu et al., 2021), which first conducted principal component analysis (PCA) on the 5 kb bin matrix. For each exposure, we selected the top 32 principal components (PCs) for clustering using the modules in scanpy (Wolf, Angerer, and Theis 2018). In the HIV-1 and influenza cohorts, we observed a donor effect in the clustering results with these PCs. Therefore, we applied harmony (Korsunsky et al., n.d.) to correct the donor effect on these PCs. We performed clustering separately for control samples (’HIV_pre’, ‘Flu_pre’, and ‘Ctrl’) and samples from the ‘MRSA/MSSA’, ‘BA’, ‘COVID-19’, and ‘OP’ groups, allowing for better comparison between the exposures and control samples.

To annotate the cells, we used both the single-cell methylation clustering results and cell surface markers. In almost every cohort, we observed two clusters of B cells and NK cells, which were distinguished by their global mCG levels. Therefore, we assigned these clusters as naive and memory B cells, naive and active NK cells. We also merged clusters with cell surface markers indicating memory CD4 and CD8 T cells, even if they exhibited multiple clusters in the T-SNE embedding.

#### Exposure-associated differentially methylated regions (eDMR) identification

To identify DMRs associated with each immune cell type, we utilized peripheral blood mononuclear cells (PBMCs) from healthy donors. Based on single-cell methylation and fluorescence-activated cell sorting (FACS), we identified nine cell types through clustering. These cell types were grouped based on their global mCG levels, and DMRs were called separately within high-mCG and low-mCG cell types. We employed methylpy (https://github.com/yupenghe/methylpy) for DMR calling, and the resulting DMRs were further annotated with genes and promoters.

To identify DMRs associated with each exposure, we merged the control samples and samples from each exposure group. We used methylpy (https://github.com/yupenghe/methylpy) to identify DMRs between the control and exposure groups and between different exposure groups. Once we obtained the primary set of DMRs, we calculated the methylation levels of all samples at these DMRs using “methylpy add-methylation-level”.

Additional filtering on the DMRs was performed by comparing the methylation levels among different sample groups using Student’s t-test. Only DMRs with a minimum p-value less than 0.05 between any two groups were retained. For DMRs associated with MRSA/MSSA, BA, OP, and SARS-CoV-2, where external controls were used for DMR calling, we compared the methylation levels of exposure samples and control samples, as well as different cohorts of controls (HIV, Flu, and commercial controls). DMRs that showed significant differences (p-value < 0.05) between the exposure group and all three control cohorts but no significant differences (p-value > 0.05) between any two control cohorts were retained.

To visualize the complex heatmaps, we employed PyComplexHeatmap (https://github.com/DingWB/PyComplexHeatmap) ^49,51^. Hypomethylated DMRs in the corresponding sample groups and cell types were labeled for better visualization. The heatmap rows were split according to sample groups, and the columns were split based on DMR groups and cell types. Within each subgroup, rows and columns were clustered using ward linkage and the Jaccard metric.

#### Validation of DMRs by shuffling the samples

To validate that the identified DMRs for each exposure were not confounded by batch effects or other factors, we shuffled the group labels of the samples within each exposure and identified DMRs among the randomly assigned groups. We quantified the methylation levels of all samples at the DMRs from the random groups and performed t-tests on the methylation levels between each pair of groups.

#### SNP Calling

We merged the bam files from the same donors, and SNP calling was performed using Biscuit’s^43^ variant calling function. This process identifies SNPs in both CpG and non-CpG contexts by analyzing the bisulfite-treated reads. Biscuit distinguishes between methylated cytosines and actual C/T polymorphisms, reducing the risk of false positives.

Standard variant filtering was applied to remove low-confidence SNPs. We excluded SNPs with a minor allele frequency (MAF) below 0.05. Additionally, SNPs overlapping with regions in the blacklist were filtered out.

#### Identification of gDMR-meQTL Pairs

To identify methylation quantitative trait loci (meQTLs) associated with differentially methylated regions (DMRs), we used QTLtools (v2.0-7-g61a04d2c5e) ^52^. The analysis was conducted using two approaches: nominal and permuted methods, both designed to account for the statistical significance of the association between SNPs and methylation levels. DMRs between the 110 donors in each cell type were identified using methylpy.

meQTL Mapping

1. Nominal Analysis: We used QTLtools’ nominal mode to calculate the association between genotype (SNP) and methylation levels within DMRs. This method tests all SNP-DMR pairs within a specified genomic window (1Mb) around the DMRs, reporting nominal p-values for each pair. Associations were considered significant at a nominal threshold of FDR < 0.01.
2. Permutation-Based Analysis:

To estimate the empirical significance of the identified meQTLs and correct for multiple testing, we performed 1000 permutations of the methylation data using QTLtools’ permutation mode. The permuted p-values were used to compute a false discovery rate (FDR) and assess the robustness of the identified DMR-meQTL pairs.

Covariates and Adjustment

In both analyses, we included covariates such as age, sex, first 5 PCs of genotypes, and exposures. Covariate adjustment was performed using the QTLtools built-in method for linear model regression.

#### Motif enrichment

We obtained the hypo- and hyper-DMRs reported by methylpy from the columns ‘hypermethylated_samples’ and ‘hypomethylated_samples’. HOMER was used to identify enriched motifs within these different sets of DMRs for each exposure. The results from HOMER’s ‘knownResults.txt’ output files were used for downstream analysis. Only motif enrichments with a p-value < 0.01 were retained. The motif enrichment results were visualized using scatterplots in seaborn.

#### Differentially methylated gene (DMG) identification

Pairwise differential methylation analysis of genes (DMGs) for each exposure was performed using ALLCools, following the tutorial (https://lhqing.github.io/ALLCools/cell_level/dmg/04-PairwiseDMG.html). Significantly differentially methylated genes were selected based on an FDR < 0.01 and a delta mCG > 0.05. The functional enrichment analysis of the DMGs was conducted using metascape (Zhou et al., 2019) (https://metascape.org/).

#### Integration with single-cell ATAC

We integrated our single-cell methylation data with single-cell ATAC-seq data from HIV-1. This integration was performed using Canonical Correlation Analysis (CCA), where we transferred our methylation cell annotations to the cells from the other modality. To generate the peaks and bigwig files for each cell type, we utilized SnapATAC2 (Zhang et al., 2021; Fang et al., 2021).

#### Correlation of single-cell methylation and single-cell ATAC

To assess the correlation between single-cell methylation and single-cell ATAC, we calculated the correlation between the hypo-score of each 5 kb bin and Tn5 insertions in each bin. This correlation was performed between cell types and within matched cell types.

#### Colocalization of meQTL with GWAS traits

Summary statistics of GWAS were downloaded from the NHGRI-EBI GWAS Catalog (https://www.ebi.ac.uk/gwas/) ^53^, including 29,401 studies and 25,111 traits. we performed colocalization analysis with coloc (v5.2.3) using default priors to calculate the probability that both the meQTL and GWAS traits share a common causal variant. The posterior probability (PP4) of one causal variant associated with both DMR and GWAS traits was used to identify significant colocalizations (PP4 > 0.50), and a high PP4 value indicates strong evidence for shared causality. R packages locuscomparer (v1.0.0)^54^ and locuszoomr (v0.3.1) ^55^ were employed to visualize the colocalization results. To test whether the meSNP (snp of meQTL) and GWAS SNP are significantly overlapped for each pair of colocalized DMR and trait, we used the chi-squared test (function stats.chi2_contingency from Python package scipy ^50^), p-values were adjusted using Benjamini/Hochberg method.

## Acknowledgments

This work was supported by the Defense Advanced Research Projects Agency (DARPA) through the DARPA Epigenetic Characterization and Observation (ECHO) program for the project Single-cell Analysis for Forensic Epigenetics (SAFE), issued by the US Army Research Office under co-operative agreement W911NF-19-2-0185. J.R.E. is an Investigator of the Howard Hughes Medical Institute. We would like to thank former DARPA Biological Technologies Offices Program Manager Eric Van Gieson, as well as the current leadership team including Jean-Paul Chretien and Thomas Thomou, for their guidance and insightful comments. We also acknowledge DARPA for contract number W911NF-19-2-0185 and Xuechen Yu. Armen Donabedian is acknowledged by TE. We extend our gratitude to the ECHO team of Arizona State University, particularly Joshua LaBaer and Vel Murugan, for coordinating the various teams during the initial phase of the study. We would like to thank all the unidentified donors who contributed biological samples for this project through our collaborators. We are thankful to Susan Kaech for consultations regarding the PBMC selection strategy, as well as Caz O’Connor and Lara Boggeman at the Salk FACS Core for their assistance in standardizing the gating strategy. W.J.G. acknowledges funding support from National Institutes of Health (NIH) grants P50-HG007735 and UM1-HG009442, UM1-HG009436. S.C.S acknowledges funding support from DARPA through grant N6600119C4022. V.G.F acknowledges funding from NIH grant 1R01AI165671. This work utilized the Stampede2 supercomputing resources at Texas Advanced Computing Center through allocation MCB130189 from the Extreme Science and Engineering Discovery Environment (XSEDE), which was supported by National Science Foundation grant number #1548562. We would like to thank Madhusudan Gujral, Lei Huang, and Remy Scott for their assistance with porting and optimization of computational tools on XSEDE resources, made possible through the XSEDE Extended Collaborative Support Service (ECSS) program. The work was conducted after Salk Institutional Review Board (IRB) approval through IRB Protocol Number: 18-0015 titled “Single Cell Analysis for Forensic Epigenetics (SAFE).” Salk Federal Wide Assurance (FWA) for the Protection of Human Subject Number: FWA00005316.

