## Supplementary Figure S1 for "Genetics and Environment Distinctively Shape the Human Immune Cell Epigenome"

### A. Gating strategy

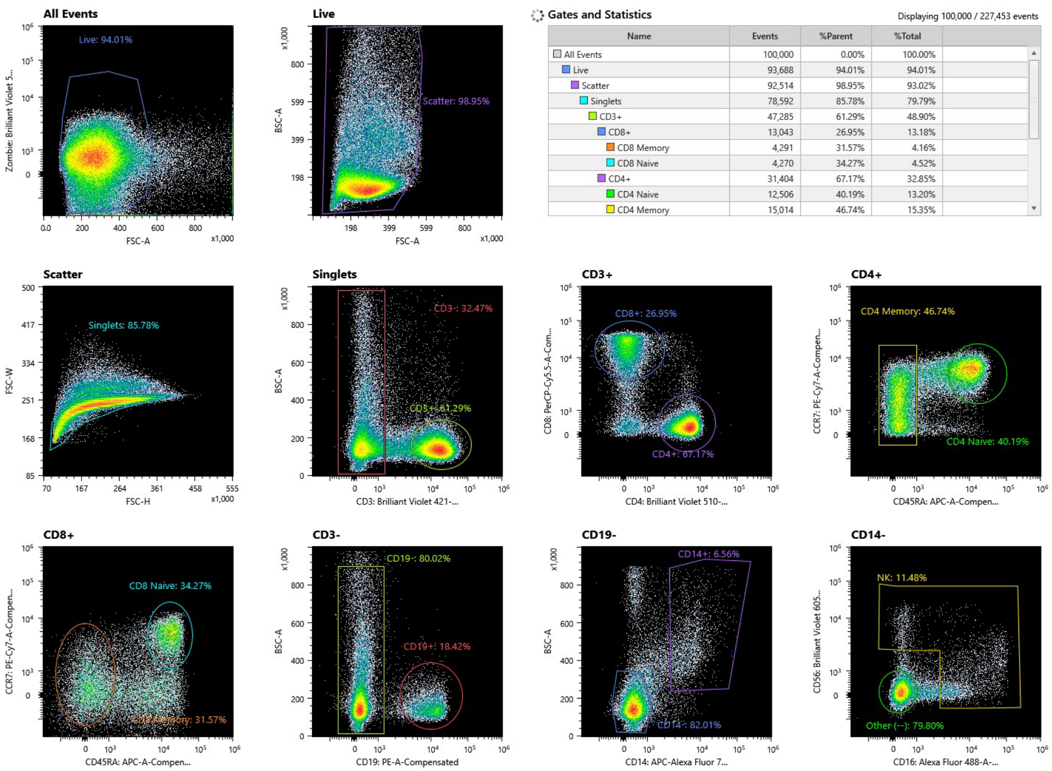


### B. Gates and Statistics from one FACS result

| Name | Events | %Parent | %Total |
|---|---|---|---|
| All Events | 100,000 | 0.00% | 100.00% |
| Live | 93,918 | 93.92% | 93.92% |
| Scatter | 92,860 | 98.87% | 92.86% |
| Singlets | 79,500 | 85.61% | 79.50% |
| CD3+ | 48,455 | 60.95% | 48.46% |
| CD8+ | 13,144 | 27.13% | 13.14% |
| CD8 Memory | 4,205 | 31.99% | 4.21% |
| CD8 Naive | 4,444 | 33.81% | 4.44% |
| CD4+ | 32,377 | 66.82% | 32.38% |
| CD4 Naive | 13,056 | 40.32% | 13.06% |
| CD4 Memory | 15,158 | 46.82% | 15.16% |
| CD3- | 25,914 | 32.60% | 25.91% |
| CD19- | 20,646 | 79.67% | 20.65% |
| CD14- | 16,736 | 81.06% | 16.74% |
| Other (--) | 13,127 | 78.44% | 13.13% |
| NK | 2,159 | 12.90% | 2.16% |
| CD14+ | 1,394 | 6.75% | 1.39% |
| CD19+ | 4,858 | 18.75% | 4.86% |

### C. Plate strategy

| Sort ID | Sort Gate | Color | Sort Mode | Cell Size | Stop Count | Timeout |
|---|---|---|---|---|---|---|
| Sort ID 1 | CD8 Memory | | Single Cell | Regular Cell | 1 | 0 |
| Sort ID 2 | NK | | Single Cell | Regular Cell | 1 | 0 |
| Sort ID 3 | CD14+ | | Single Cell | Regular Cell | 1 | 0 |
| Sort ID 4 | CD19+ | | Single Cell | Regular Cell | 1 | 0 |
| Sort ID 5 | CD4 Memory | | Single Cell | Regular Cell | 1 | 0 |
| Sort ID 6 | CD4 Naive | | Single Cell | Regular Cell | 1 | 0 |
| Sort ID 7 | CD8 Memory | | Single Cell | Regular Cell | 1 | 0 |
| Sort ID 8 | CD8 Naive | | Single Cell | Regular Cell | 1 | 0 |

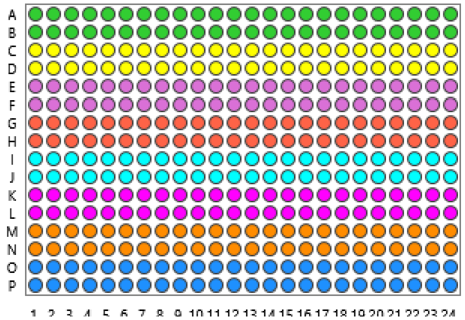
