## Supplementary figures and images for "Genetics and Environment Distinctively Shape the Human Immune Cell Epigenome"

### Supplementary Figure S2

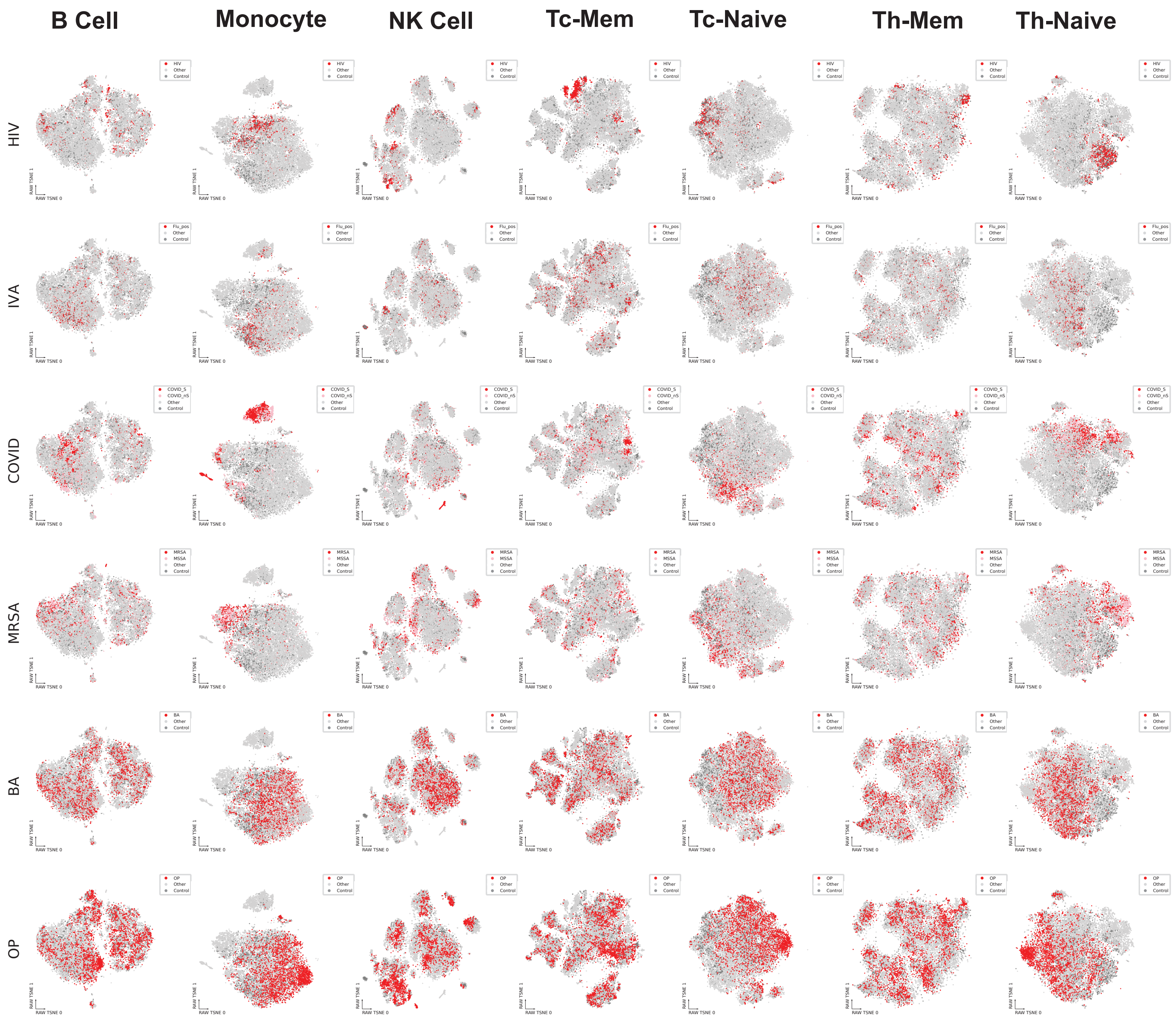

### Supplementary Figure S3

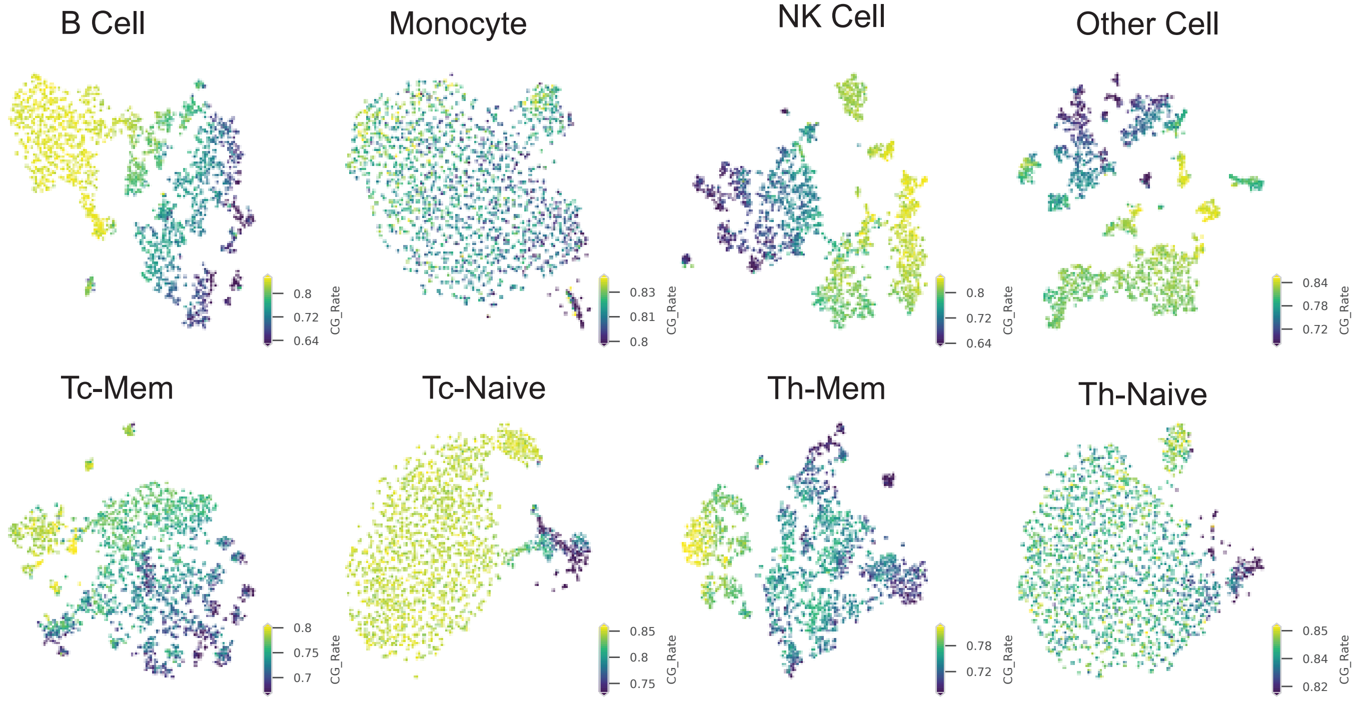

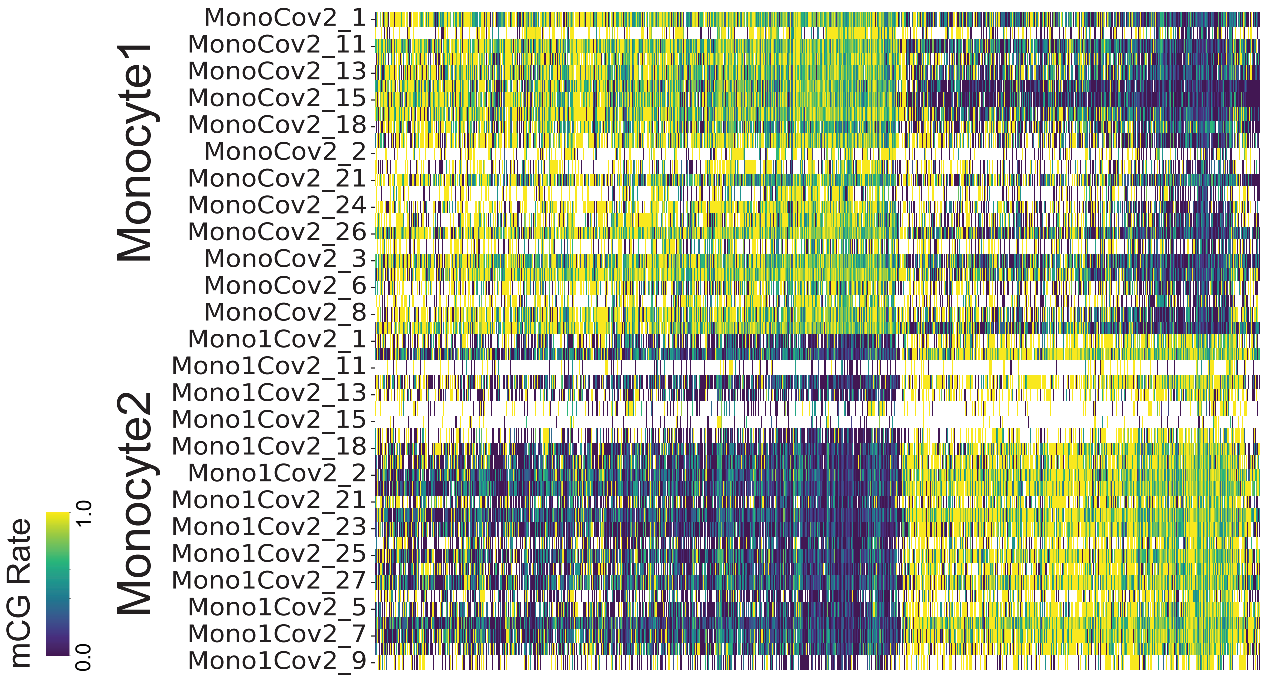

### Supplementary Figure S4

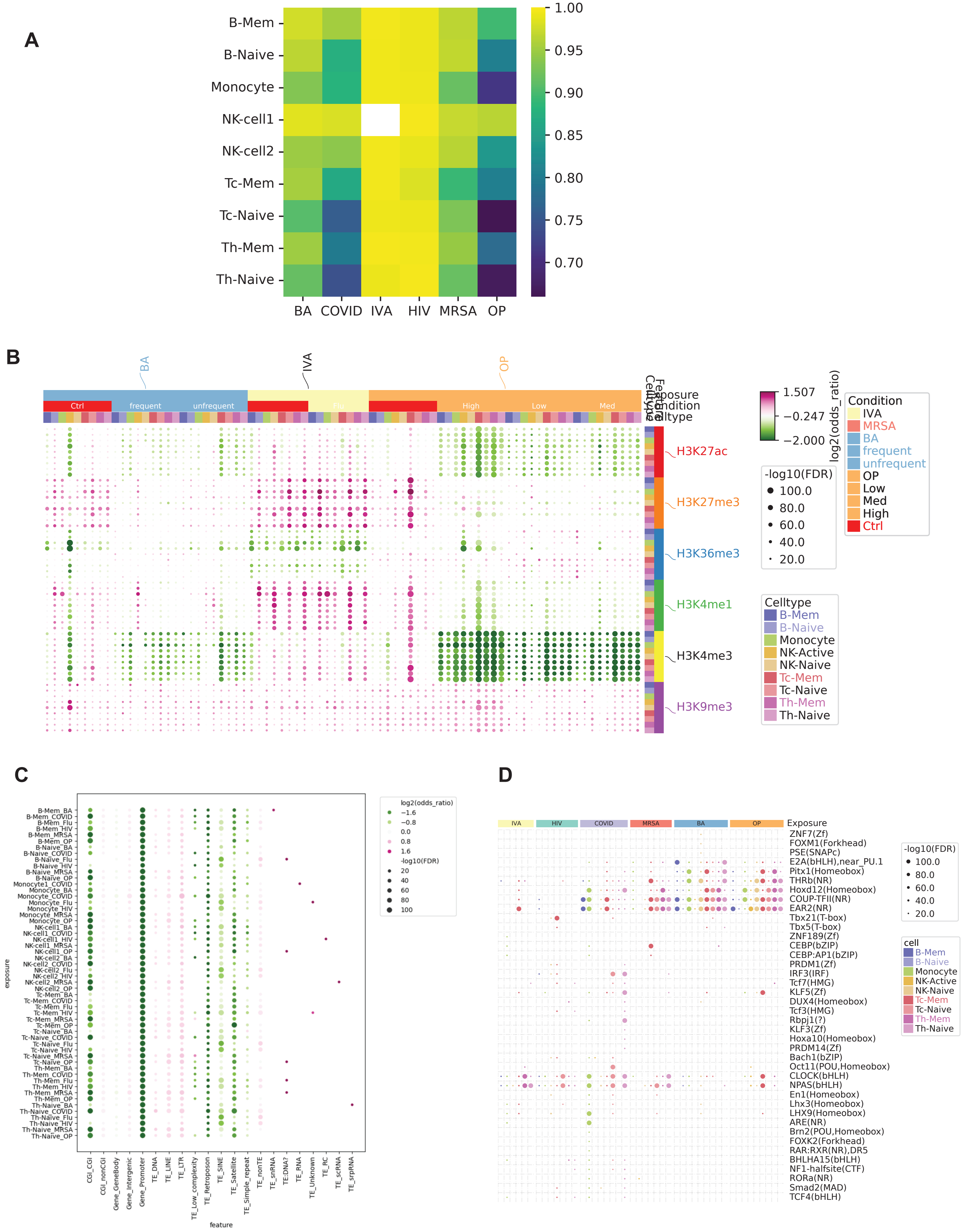

### Supplementary Figure S5

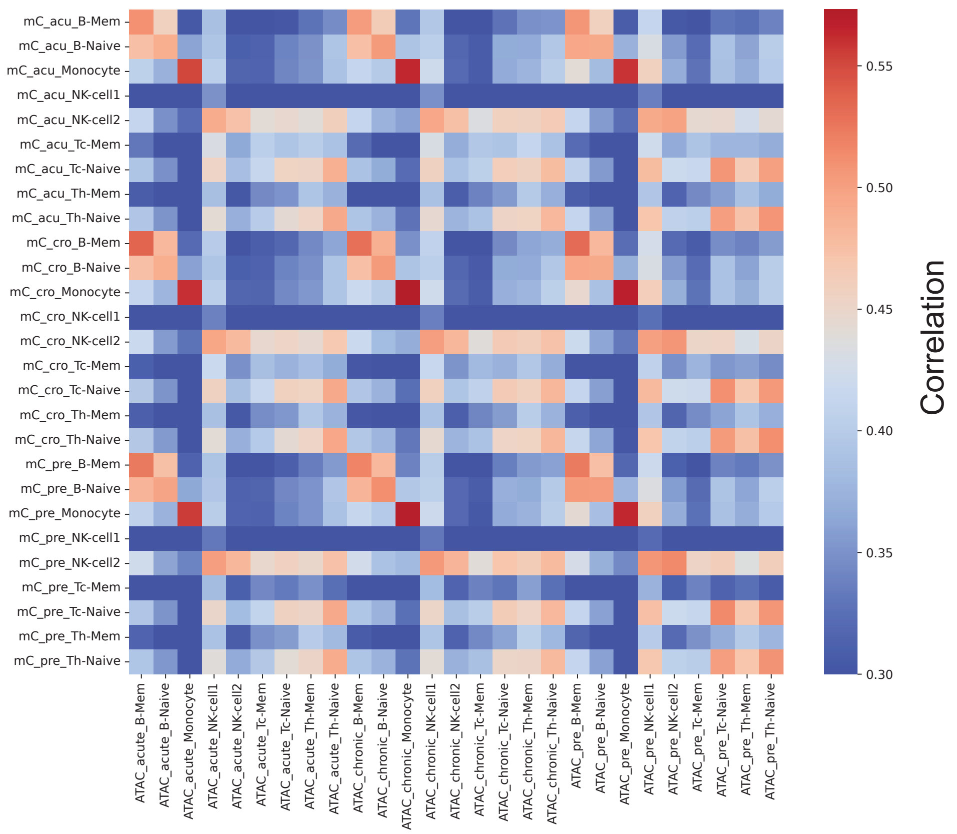

### Supplementary Figure S6

#### Figure S6

A

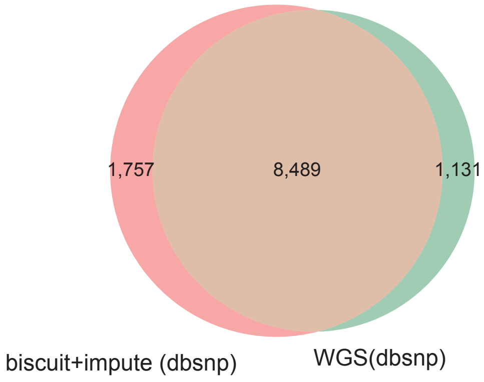


B

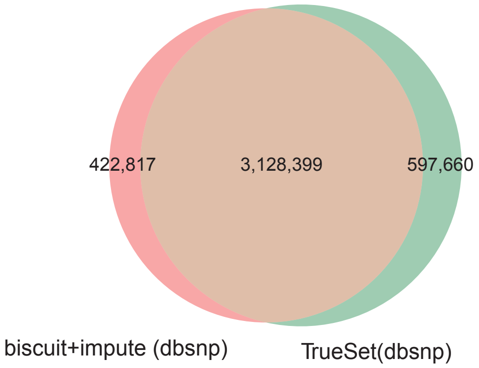


C

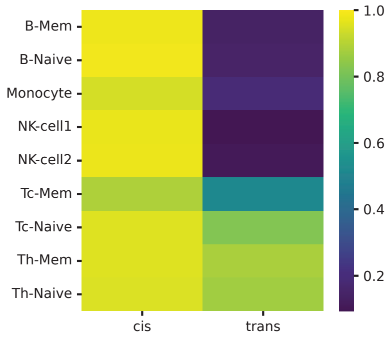


D

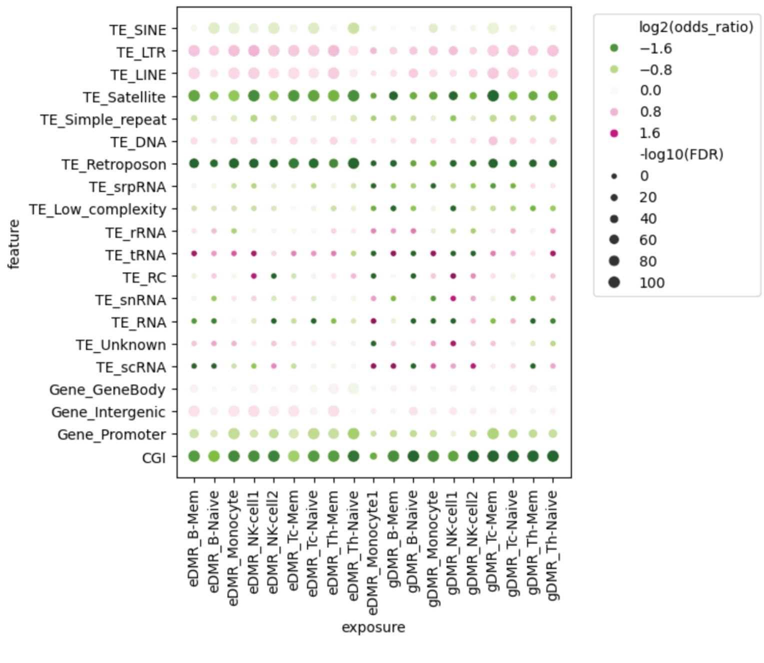


E

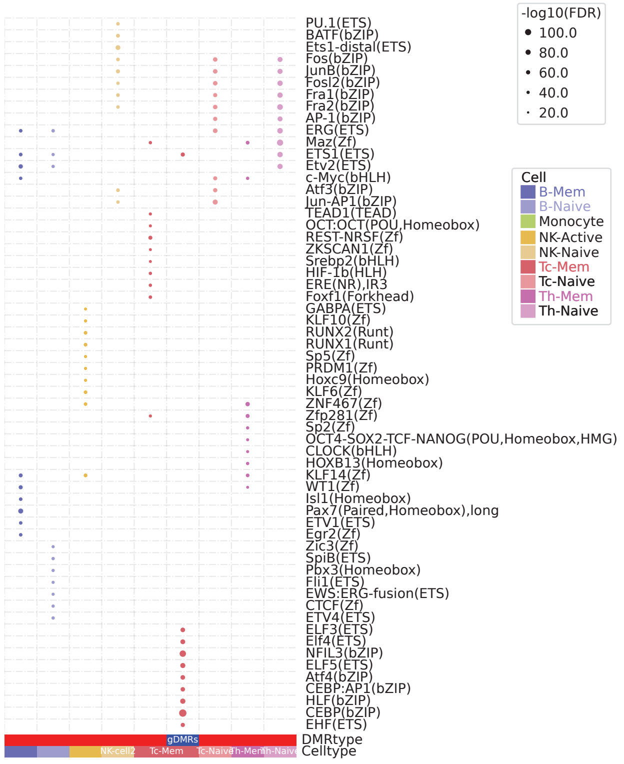

### Supplementary Figure S7

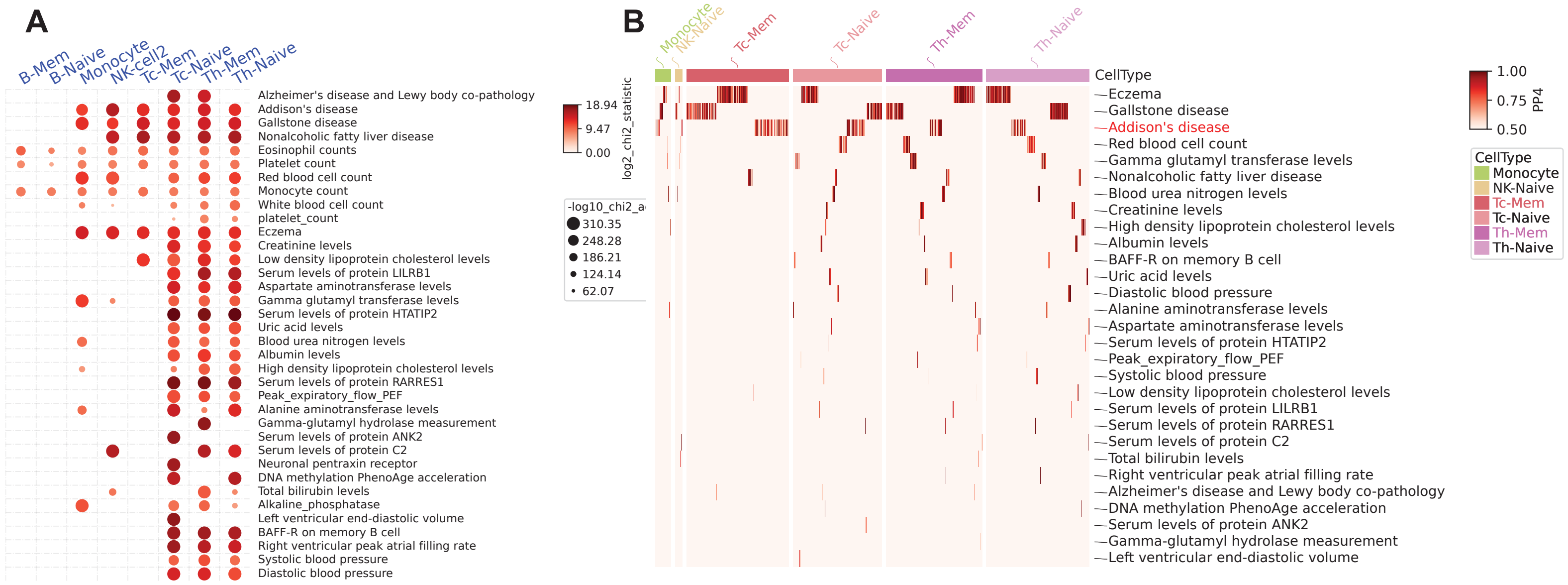
